# Non-canonical alternate interactors of SARS CoV-2 Spike protein implicated in placental susceptibilty

**DOI:** 10.64898/2026.10.08.757814

**Authors:** Sulagna Mandal, Satyabrata Majumder, Suman Kumar Nandy, Ruchira Mukherjee, Kalyan Giri, Aparna Mukhopadhyay

**Author notes:** SM is currently located in Department of Physics, University of Vermont, USA. Corresponding author: Dr. Aparna Mukhopadhyay, Molecular Virology Laboratory, Department of Life Sciences, 86/1 College Street, Presidency University, Kolkata-700073, India. This work is part of a thesis to be submitted in partial fulfilment of the requirements of the degree of Doctor of Philosophy from Presidency University, Kolkata.

## Abstract

Vertical transmission of SARS-CoV-2 is controversial. Owing to the low expression of ACE2 along with TMPRSS2 in placenta, role of secondary receptors becomes significant. In our study we predicted twelve interactors of spike protein expressed in placenta through interactome analysis followed by docking and molecular dynamic simulation. Three interactors of SARS-CoV-2 spike protein were found to be CD147, EGFR and LDLR. Expression at transcriptomic and translational level by real-time PCR and immunohistochemistry respectively was observed in placenta samples obtained from women with COVID exposure risk and compared to control. Elevated protein levels of CD147 (OR=14.667), ACE2 (OR=8.25) and LDLR (OR=1.9) was associated with higher exposure risk. However, the variation in mRNA expression was not statistically significant. Computational study showed that CD147/basigin, EGFR and LDLR stably interact with spike protein outside its primary receptor binding domain. In-vitro binding assay confirmed the physical interaction between spike protein and the placental receptors. In-silico site-directed mutagenesis further identified key potential binding hotspots of spike protein-Tyr28 and Trp64 for CD147, Phe643, Val642, Thr632 for LDLR and Leu8, Phe59, Phe32, Tyr28, Trp64, Trp152, His245, and Val213 for EGFR. Finally, concluded that CD147, EGFR and LDLR might act as secondary non-canonical cofactors for SARS-CoV-2 spike protein in placenta, thereby increasing its susceptibility and tissue tropism. The study overall gives a better insight into potential placental suscpetibility providing targeted therapeutic opportunities.

## Introduction

Since the dawn of COVID-19, vertical transmission has been a conflicting area of research. To understand the depth of vertical transmission, a more detailed understanding of molecular mechanisms involved is required. Molecular mechanisms rely on the cellular receptors as seen in other viruses like HIV, ZIKA and other vertically transmitted viruses. Although, ACE2 is the primary receptor of the SARS CoV-2 surface protein spike for viral entry in the target organ, lung, existing literature debates over its expression in the placenta. While its expression is lower in placenta than other vital organs such as lung, intestine, a few studies claim otherwise(1,2). However, most of the current literature point towards a low transcriptomic level of ACE2 in the placenta(2). Studies further suggest a gradual decrease in the expression level with advancing gestation. Furthermore, TMPRSS2, the other important cofactor of SARS-CoV-2 entry, is absent or only minimally expressed in the placenta (1–8). Therefore, the role of other alternate receptors needs to be thoroughly investigated.

The aim of this study was to characterize the placental receptors initially screened through interactome analysis by text mining existing literature (9). Among 12 possible interactors screened, three receptors-CD147/basigin, EGFR/Epidermal Growth Factor and LDLR/Low density lipoprotein receptor were chosen for the present study based on structure availability, their role in entry with respect to other viruses and literature availability. However, the other receptors might also have important roles and hence holds potential for future studies.

CD147 or basigin is a highly glycosylated type-1 transmembrane protein which acts as an extracellular matrix metalloproteinase inducer under normal physiological as well as pathological conditions. Among the four isoforms, basigin 2 is the most abundantly expressed protein (10). Expressed in blastocysts and term placenta, basigin plays important role in regulating trophoblast functions (11,12). Studies suggest that HIV infection is enhanced by its interaction with CD147 and cyclophilin A (13). Besides, its role in SARS-CoV-2 entry has been implicated since the beginning of the pandemic. There are studies that support the notion that SARS-CoV-2 infection might alter the expression of CD147 in placenta (14). Another study suggested the role of CD147 in COVID infection susceptibility of digestive tracts which was in turn associated with high VEGF level (15).

Another transmembrane protein epidermal growth factor receptor or EGFR is important for cell proliferation, growth, development and survival. It has the highest expression in placenta compared to other human non-malignant tissues and plays critical role in placental development and survival (16,17). It is known to protect the placental trophoblasts from apoptosis (18). There are also viruses like Zika, HCV, HBV, EBV that exploit EGFR for their entry through endocytosis (19). Similarly, their role in COVID infection has also been suggested by a few studies. A cross-talk between ACE2 and EGFR has been suggested where ACE2 is shown to negatively regulate EGFR-Ras GTPase activation and an EGFR-dependent activation of MAPK is demonstrated by spike-RBD induction (20). Furthermore, EGFR was found to be overexpressed in lung tissues of COVID-19 deceased patients and use of nimotuzumab, a EGFR specific antibody could prevent fibrosis in severe and moderate COVID-19 patients (21). Association of EGFR with severity of pulmonary fibrosis with elevated CPR levels in COVID patients was also suggested by another study (22). The possible hijacking of EGFR signalling pathways by SARS-CoV-2 spike protein has been reported in A549 lung cancer cells. Furthermore, AKT activation induced by spike-RBD was demonstrated to happen in a EGFR-dependent manner (23).

Our third target receptor LDLR or low-density lipoprotein receptor is another glycoprotein which plays a very symbolic role in the synthesis and transport of cholesterol in humans. It has the highest expression in the placenta during onset of pregnancy (24). The receptor number is regulated by its cellular demand for cholesterol on which progesterone synthesis is primarily dependent (25). What makes it an interesting candidate is its role in viral entry for HCV and vesicular stomatitis virus (26,27). It also acts as a host factor for flaviviral entry and infection in neurons (28). Similarly, its role in COVID-19 infection has also been under intense research. Severe COVID-19 patients have been shown to downregulate LDLR signalling pathway which is associated with poor disease prognosis (29). Also SARS-CoV-2 pseudovirion infection of retinal ARPE-19 cells was confirmed to be mediated by LDLR (30). Very recently, LDLR has been identified as a driver of COVID-19 myocarditis through its interaction with osteopontin (OPN) (31).

In this study, we demonstrate the protein and RNA levels of the selected receptors i.e. CD147, EGFR and LDLR in placental samples, study their interaction *in silico* and follow up with *in vitro* binding studies. *In silico* mutagenesis studies further indicate their potential to be established as novel cofactors for SARS CoV-2 entry.

## Methods

### Interactome analysis

The interactome analysis was done by listing all the proteins expressed in the placenta tissue from Human Protein Atlas (32), UniProt (33) and an extensive text mining for studies on placenta proteins as well as studies on placenta protein interaction with structural and non-structural proteins of SARS-CoV-2 (9,34,35). Only membrane proteins expressed in the placenta confirmed by tissue specificity through UniProt database was considered in the current study. The topological domains of the screened proteins was determined and compared from UniProt, TMHMM (36) and Phobius (37) to locate the extracellular domain which is hypothesised to bind with the ecto domain of viral spike protein. Expression of the proteins were further confirmed by TISSUES (38) and TiGER (39) web resources. A comparative expression level of these proteins in the placenta was also checked using C-IT (40) and BioGPS (41) databases based on Z-score value and median score respectively. After determining the above properties, interaction between the screened placental proteins and SARS-CoV-2 spike protein was predicted using VirusHostPPI (42), HVPPI (43)and UniProt databases. Finally, 12 placental membrane interactors of spike protein were listed with at least two positive hits from the three above-mentioned databases.

### Sample collection

Placenta tissue collection was done after obtaining the human ethical clearance from Calcutta Medical College Hospital, Kolkata and Calcutta National Medical College Hospital, Kolkata (vide ethical certificate: MC/KOL/IEC/NON_SPON/1290/03/2022 dated 16/03/2022 and EC-CNMC/2024/403 dated 04/05/2024). Sample collection has been detailed in our previous publication (44). Briefly, total 30 control and 30 placenta samples from women determined to be at risk of COVID exposure were collected. Risk of COVID exposure was determined by a questionnaire and the risk was determined by a scoring mechanism detailed in the paper (44) Due to limitations of biosafety protocols, placenta was collected from COVID negative patients at the time of delivery only. The placental tissue was obtained after vertical incision, 2 centimetres away from the umbilical cord to collect the chorionic villi region as it was our area of interest. Tissues were divided into two parts for downstream experiments. One was fixed in 10% formalin solution while the other part was stored at -80°C.

### Immunohistochemistry of placenta tissues and quantification

Tissue blocks were made using paraffin wax and 4-micron tissue sections were cut using Leica microtome. The tissue sections were fixed on charged slides and deparaffinized using xylene. Gradual rehydration steps of 100%, 90%, 70% and 50% ethanol were followed before antigen retrieval using citrate buffer at 85°C for 45 minutes. To quench the endogenous peroxidase activity, tissue sections were treated with 3% H_2O2 f_or 20 minutes in dark. Sections were washed with 1X Phosphate Buffer Saline (PBS) and blocked with 5% Bovine Serum Albumin (BSA) at room temperature for 1 hour. This was followed by incubation with primary antibodies-CD147 Rabbit pAb, EGFR Rabbit pAb, LDLR Rabbit pAb and ACE2 Rabbit pAb (ABclonal, 1:200) overnight at 4°C. Next day sections were washed with 1X PBS twice before incubation with secondary antibodies at room temperature for 1 hour. Finally, sections were developed using 3,3’-diaminobenzidine (DAB) (Santa Cruz Biotechnology) for 1 hour at RT in dark and counterstained with haematoxylin. Post haematoxylin, gradual dehydration was done using different concentrations-50%, 70%, 90% of alcohol for 5 minutes each. Slides were cleared in Xylene for 10 minutes and mounted with DPX. Images were captured under 20X magnification by Leica DMi 8 microscope using LASX software.

To semi-quantitatively assess the receptors’ expression level in the placenta tissue, a scoring system described previously was used (45,46). The total score was a sum of (i) percentage of stained villi and (ii) staining intensity. Out of a total of 100 chorionic villi per sample, if the DAB positive villi were between 1-20, a scoring of 1 was assigned whereas scores of 2,3 and 4 were recorded for 21-40, 41-60 and >60 DAB positive villi respectively. Intensity was scored as 1, 2 and 3 based on weak, moderate and strong staining. The points were added to give a final scoring which would semi-quantitatively determine the expression level of the receptors as low, medium and high. To remove bias, the scoring was independently done by two investigators.

### qPCR analysis

Total mRNA was isolated from the placenta tissues. They were homogenized in Trizol, followed by phase separation by addition of 100uL of chloroform for 500uL Trizol. RNA was precipitated by isopropanol and washed with 75% ethanol. The pellet was air-dried and dissolved in DEPC treated water at 65°C for 5 minutes. The aliquots were stored at -80°C. The cDNA was made with 185ng/uL of RNA using iScript Reverse Transcriptase Supermix (BioRad). The qPCR reactions were performed in duplicates for each receptor using primers specific for ACE2, CD147, EGFR and LDLR. SSOFAST Evagreen Supermix (BioRad) was used and the reactions were run on BioRad CFX Duet Real-Time PCR System using BioRad CFX Maestro Software 2.3. Finally, fold change expression (-ΔΔCq) of each receptor was calculated after normalization with GAPDH expression and analysed statistically.

### Protein-protein docking

Protein sequences were sourced from the UniProt database, and their corresponding three-dimensional structures for molecular docking and molecular dynamics (MD) simulations were retrieved from the Protein Data Bank (PDB). The relevant PDB and UniProt IDs for the proteins analyzed in this study are as follows: SARS-CoV-2 Delta variant spike protein (PDB ID: 7W92 (46), UniProt ID: P0DTC2), Basigin (PDB ID: 4U0Q (47), UniProt ID: P35613), Epidermal Growth Factor Receptor (EGFR) (PDB IDs: 1NQL (48), 3NJP (49), 3QWQ (50), 5XWD (51), 1IVO (52), UniProt ID: P00533), Low-Density Lipoprotein Receptor (LDLR) (PDB ID: 3M0C, UniProt ID: P01130). Missing residues in the protein structures were modelled using the CHARMM-GUI (53) server, while the protonation states of residues at pH 7.4 were assigned using the PROPKA (54) tool.

We used ClusPro (55), a rigid-body docking server, to perform blind docking of the Spike protein with EGFR and LDLR, and CD147/basigin. The server executes three computational steps: (1) rigid body docking, sampling billions of conformations; (2) clustering of the 1,000 lowest energy structures based on root-mean-square deviation (RMSD) to identify the largest clusters representing the most probable models of the complex; and (3) refinement of selected structures through Charmm energy minimization. The rigid body docking phase utilizes PIPER (56), a docking program based on the Fast Fourier Transform (FFT) correlation method. Through benchmark-based evaluations and an overview of the CAPRI and CAPRI/CASP experiments, it was observed that ClusPro generates a higher number of high-quality models among the top 5 and top 10 predictions compared to the best flexible docking methods (57).

### Molecular Dynamics (MD) simulation and post simulation analysis

The top-scoring models derived from the docking calculations were subjected to MD simulations to assess their conformational dynamics and stability. The AMBER99SB-ILDN force field (58) and TIP3P (59) water model were used. Each protein complex was solvated in a cubic water box with a minimum protein-to-box distance of 15 Å, and Na⁺ and Cl⁻ ions were added to neutralize the system. Energy minimization was performed using the steepest-descent algorithm for up to 50,000 steps. The systems were then equilibrated for 100 ps under the NVT ensemble, with the temperature gradually increased to 300 K, followed by 100 ps of NPT equilibration at 300 K and 1 atm. During both equilibration phases, protein heavy atoms were position-restrained with a force constant of 1000 kJ mol⁻¹ nm⁻². A 2 fs time step was used throughout, followed by an unrestrained 20 ns or 50 ns production simulation. Long-range electrostatic interactions were treated using the smooth particle-mesh Ewald method (60), with a 12 Å real-space cutoff; the same cutoff was applied to van der Waals interactions. Residue–residue interaction frequencies were calculated using GetContacts (https://getcontacts.github.io/static_to_fingerprints.html). Additional trajectory metrics were obtained using built-in GROMACS tools. Data were visualized using Python, and molecular structures were rendered using PyMOL. All the simulations were performed using GROMACS 2021 (61) with GPU acceleration.

### Cell lysate and tissue lysate preparation

For coimmunoprecipitation experiments, cell lysate expressing spike protein was prepared form 293T cells transfected with spike construct obtained from Dr. Marc C. Johnson (47). Transfection was done using TurboFect Transfection Reagent (Thermo Fisher Scientific, Lithuania) using manufacturer’s instructions. 48 hours post transfection cell lysates were prepared. Confluent cells were washed and collected, followed by addition of 100uL of lysis buffer (50mM Tris (pH7.6), 150mM NaCl, 5mM MgCl_2,_ 0.5% Triton X, protease inhibitor cocktail, protease inhibitor powder, phenylmethylsulfonyl fluoride (PMSF)). Following an incubation period of 1 hour on ice, they were centrifuged at 14000rcf for 20 minutes and lysates were collected and stored at -20°C.

Coimmunoprecipitation experiments required placenta tissue lysates expressing the receptors-CD147, EGFR and LDLR. Previously stored placenta tissues from -80°C freezer were weighed and homogenized using 1.5mL of Radioimmunoprecipitation (RIPA) buffer (150mM NaCl, 1% NP-40, 0.5% sodium deoxycholate, 0.1% SDS, 50mM Tris pH 8). The homogenate was centrifuged at 13,000xg for 15 minutes at 4°C. The lysate was collected, aliquoted and stored at -80°C.

### Coimmunoprecipitation of spike with CD147, EGFR and LDLR

To validate the *in-silico* interactions predicted, coimmunoprecipitation was performed to confirm the binding *in-vitro*. A slurry of 40uL protein AG agarose beads (BioBharati LifeScience Pvt.Ltd.) was taken and blocked by addition of 5% bovine serum albumin to inhibit any non-specific interaction. The protein of interest, CD147 was immobilised on agarose beads using 1.52ug of CD147 Rabbit pAb antibody (ABclonal) which was incubated with 70uL placental lysate for 2 hours in a rotor at 20 rpm at 4°C. Control sets did not have any antibody. This was followed by washing the beads 5 times with lysis buffer (50mM Tris (pH7.6), 150mM NaCl, 5mM MgCl_2,_ 0.5% Triton X). Interaction with spike was assessed by incubation with 85uL of spike protein expressing 293T cell lysate for 2 hours in a rotor at 20rpm at 4°C followed by collection of the unbound lysates. The bound lysate was collected after washing the beads 5 times with lysis buffer and stored at -20°C. The same protocol was followed for EGFR and LDLR as well using EGFR Rabbit pAb and LDLR Rabbit pAb antibody respectively. Each coimmunoprecipitation experiment was repeated 3 times.

### Western blot

After performing the coimmunoprecipitation experiments, the binding was determined by western blot. Immunoblotting was done to observe the presence of spike protein which interacted with the placental receptors. Therefore, bound protein and unbound protein lysates along with cell lysate expressing spike protein were separated by 10% SDS-PAGE. The proteins were then transferred on Polyvinylidene fluoride (PVDF) membrane for 1.5 hours at 110 Volts. The transfer was followed by blocking the membrane with 5% skimmed milk for 30 minutes. The membrane was subsequently incubated with SARS-CoV-2 Spike Rabbit pAb primary antibody (1:1000) overnight at 4°C. Next day the membrane was washed 3 times with 1X TBST followed by incubation in HRP-conjugated anti-rabbit secondary antibody (1:5000) (BioBharati LifeScience Pvt.Ltd.) for 1 hour at 4°C. Post incubation, the membrane was washed 3 times with 1X TBST and developed with enhanced chemiluminescent substrate (ECL) (ABclonal). Finally, the blot was captured with Chemidoc (BioRad laboratories). For all 3 receptors the same protocol was repeated.

### *In-silico* site directed mutagenesis

An *in-silico* saturation mutagenesis was performed for the spike-basigin, spike-LDLR and spike-EGFR complexes to confirm the role of the interacting residues. BeAtMuSiC (62) was used to carry out the saturation mutagenesis in the interfacial residues between the target and receptor proteins. In all cases, all the chains of the target protein were considered while running the program. The effect of single-site mutations on protein–protein binding affinity was evaluated as the change in binding free energy (ΔΔG_bind). Positive ΔΔG_bind values indicate mutations predicted to weaken binding, whereas negative values indicate substitutions predicted to strengthen binding relative to the corresponding wild-type residues.

### Statistical analysis

All data was analysed using IBM SPSS v26 and MS excel. Odds ratio analysis with 95% confidence interval was performed to assess the associations. The association-based observations were further reiterated using Pearson chi-square (ꭓ^2^) analysis. The results have been tabulated and graphically represented following standard norms.

## Results

### Interactome Analysis identified twelve possible interactors of Spike protein in placenta

Through interactome analysis twelve possible interactors of spike protein were identified in placenta (**Table 1**). As detailed in methods section, they were shortlisted from the extensive screening process through collection and comparison of placental gene expression data, recognition of extra cellular domains and prediction of host virus interaction as depicted in the supplementary document Interactome analysis workflow.xlsx.

**Table 1:** Interactors of spike protein in placenta via interactome analysis.

| Protein name | Gene name | UniProt ID |
| --- | --- | --- |
| Angiotensin-converting enzyme 2 | ACE2 | Q9BYF1 |
| C-type lectin domain family 4 member M | CLEC4M | Q9H2X3 |
| Basigin | BSG | P35613 |
| Epidermal growth factor receptor | EGFR | P00533 |
| Neuropilin-1 | NRP1 | O14786 |
| CD209 antigen | CD209 | Q9NNX6 |
| Integrin beta-1 | ITGB1 | P05556 |
| Low-density lipoprotein receptor | LDLR | P01130 |
| E3 ubiquitin-protein ligase NEDD4 | NEDD4 | P46934 |
| Syndecan-4 | SDC4 | P31431 |
| Transient receptor potential cation channel subfamily V member 2 | TRPV2 | Q9Y5S1 |
| Amyloid-beta precursor protein | APP | P05067 |

### CD147/basigin, EGFR and LDLR are expressed in the placenta tissue

After completion of the interactome analysis, three candidate proteins – CD147, EGFR and LDLR were chosen to initiate the downstream experiments based on structure availability, their role in viral entry in general and availability of literature. Before proceeding to check their expression at the transcriptional and translational level in patient samples, their expression in placenta tissue was checked by western blot and distinct bands were observed for all three receptors (**Figure 1**).

**Figure 1.**
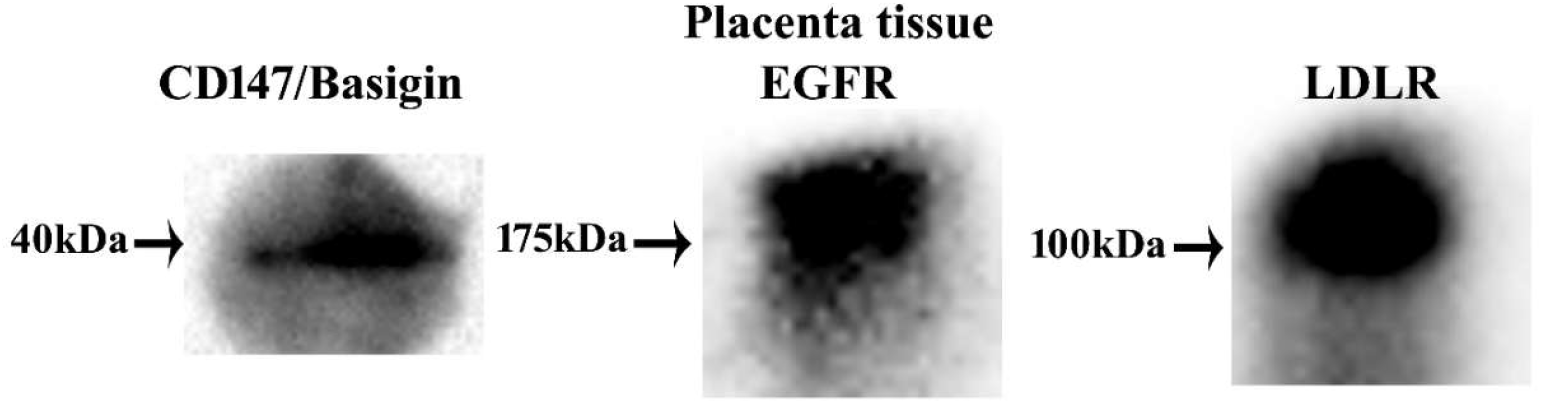
Expression of CD147/basigin, EGFR and LDLR in placenta tissue by western blot: Expression of candidate proteins – CD147, EGFR and LDLR in the placenta tissue. Distinct bands were observed for all three receptors. The observed band size for Basigin, EGFR and LDLR are 40kDa, 175kDa and 100kDa respectively.

### Change in mRNA and protein expression of the receptors is associated with COVID exposure-risk

To assess whether the three receptors under consideration differ in placental tissue with COVID exposure risk, we obtained placental samples from women in accordance to clearance obtained from Calcutta Medical College and National Medical College. As detailed in our earlier publication (44), COVID exposure risk was assessed in the women via questionnaire and categorised as per the score developed. Transcription level expression was assessed by reverse-transcriptase PCR whereas protein level expression was observed via immunohistochemistry and scored as detailed in the methods section. As the interface for viral transmission from mother to fetus is via the trophoblast, it was important to assess the expression of the proteins specifically in the trophoblast region. Hence immunohistochemistry was preferred over immunoblotting. When compared between the control patients and those at risk of being exposed to COVID during their pregnancy, it was found that there was a minor, statistically insignificant difference in the expression level (both at mRNA and protein level) of our predicted receptors. For both CD147/basigin and LDLR, expression was relatively higher in women at risk compared to the control group at the transcriptional level as well as protein level (**Figure 2**, **3(B), 3(D)**). Although, the EGFR protein expression was slightly higher in exposed-risk group than the control group, its corresponding mRNA expression was low in the pregnant women with epidemiological exposure risk compared to the control women. Furthermore, ACE2, the canonical receptor of SARS-CoV-2, was found to have a higher mRNA and protein expression in the patients with higher risk of COVID exposure than the control group (**Figure 2**, **3(C), 3(A)** respectively). Odds ratio values (**Table 2 and 3**) indicate positive association between epidemiological exposure risk and receptor expression, both at mRNA and protein level for all the receptors with exception for EGFR. Although, transcriptional changes of the interactors did not reach statistical significance (p>0.05), positive trends in protein expression, except EGFR whose protein expression remains unchanged and corresponding mRNA level shows lower expression with epidemiological exposure risk, indicate that maternal environmental risk factors may potentially modulate placental microenvironment, rendering them susceptible for viral entry.

**Figure 2:**
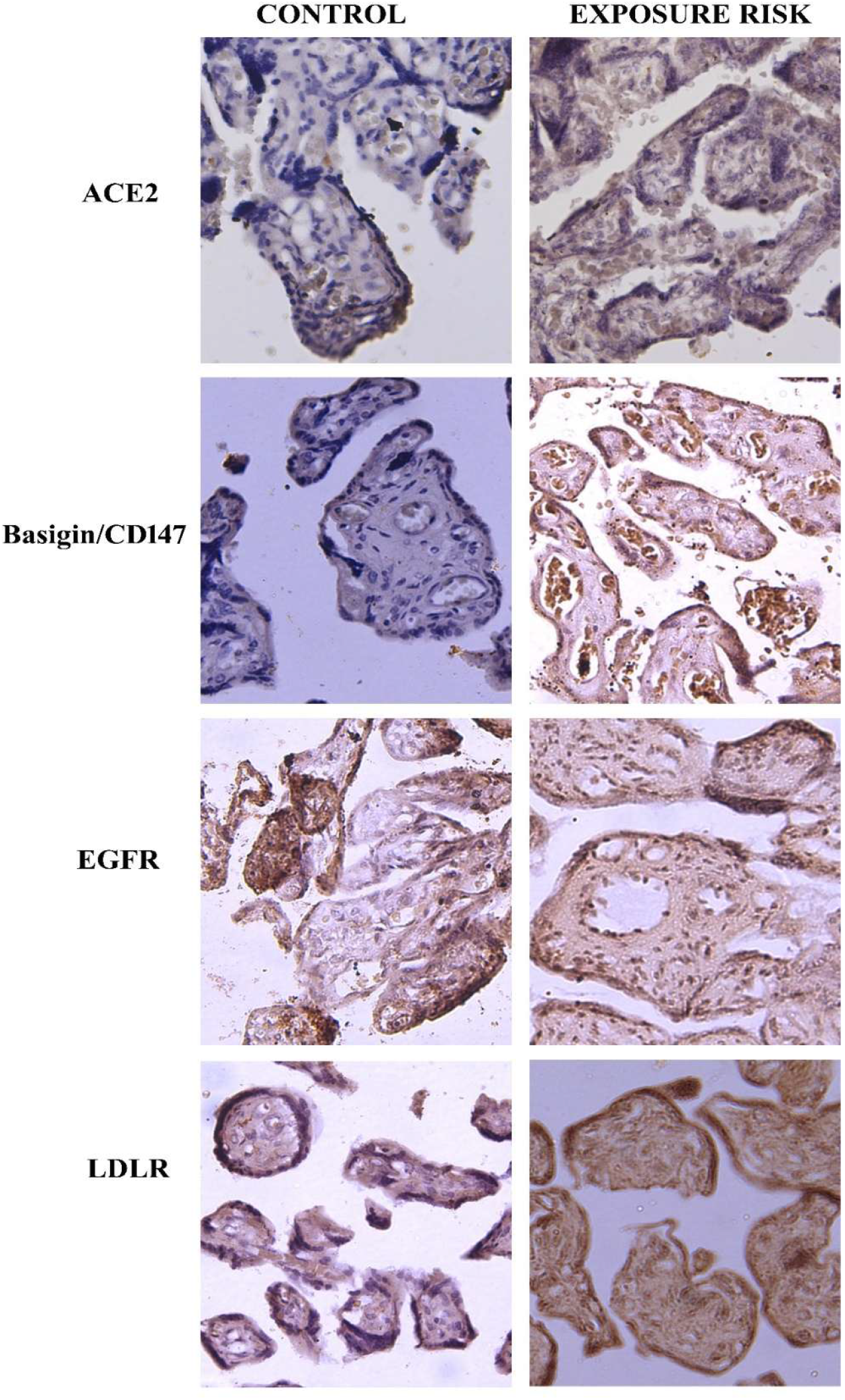
Expression level of receptor proteins in placenta by immunohistochemistry: Representative figures of protein expression and localisation of ACE2, CD147/basigin, EGFR and LDLR as determined by immunohistochemistry. The left panel shows the protein expression level in control group and the right panel shows receptor protein expression changes in pregnant women with high exposure risk to SARS-CoV-2.

**Figure 3:**
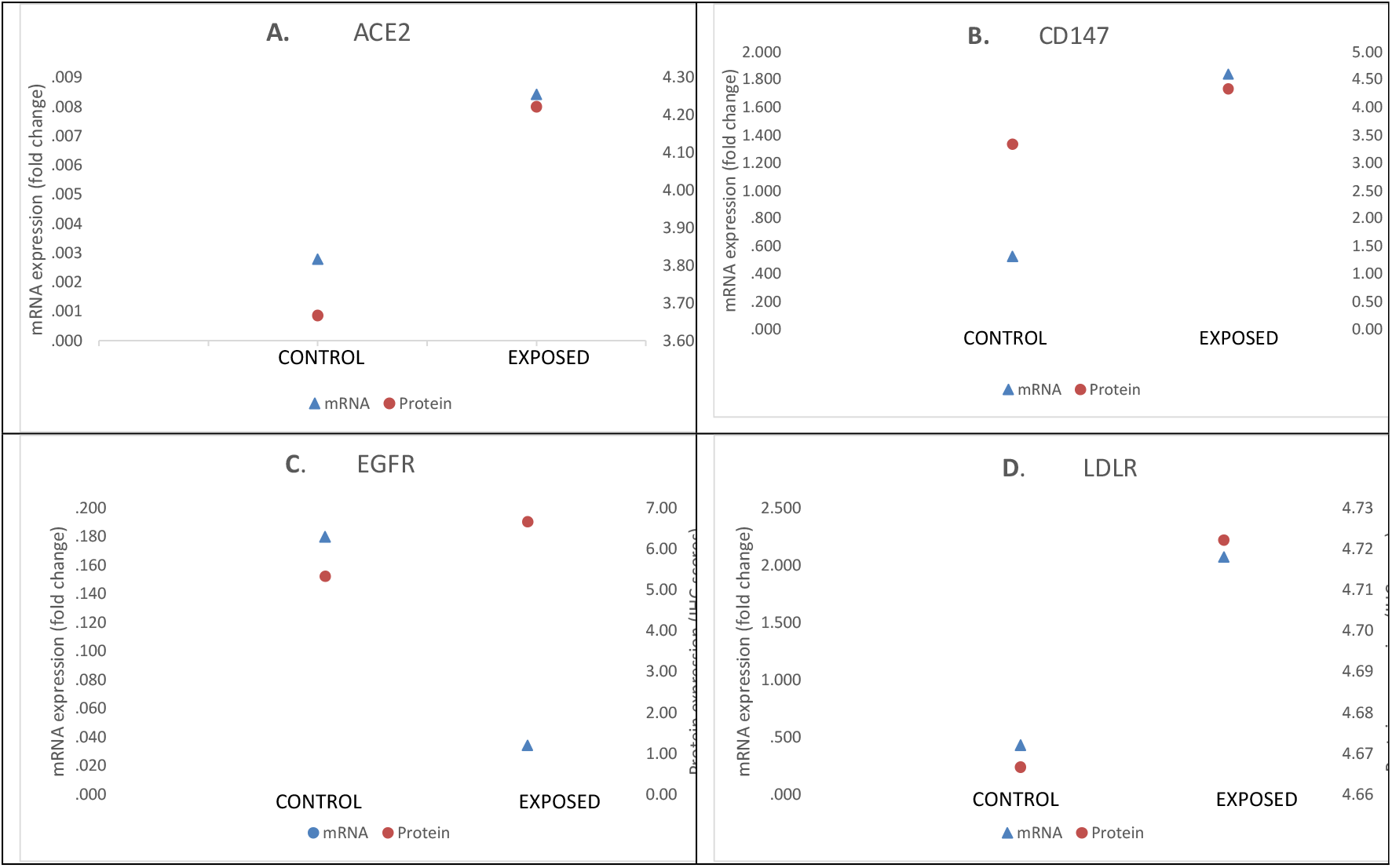
Protein expression (by quantification of IHC) and mRNA expression (by qRT-PCR) of (A) ACE2 (B) CD147/basigin (C) EGFR (D) LDLR in placental samples obtained from control women and those with exposure risk to COVID during pregnancy. Blue triangles indicate mRNA expression; red dots indicate the corresponding protein expression.

**Table 2:** The association of receptor mRNA expression to COVID exposure risk obtained by real-time PCR.

| Receptor | Odds ratio | Lower limit | Upper limit | Pearson chi-square ($\chi^2$) | p-value |
| --- | --- | --- | --- | --- | --- |
| CD147/basigin | 2.22 | 0.334 | 14.803 | 0.697 | 0.404(p>0.05) |
| LDLR | 6.667 | 0.641 | 69.344 | 3.022 | 0.082(p>0.05) |
| EGFR | 0.692 | 0.128 | 3.752 | 0.183 | 0.669(p>0.05) |
| ACE2 | 1.083 | 0.182 | 6.439 | 0.008 | 0.930(p>0.05) |

**Table 3:**
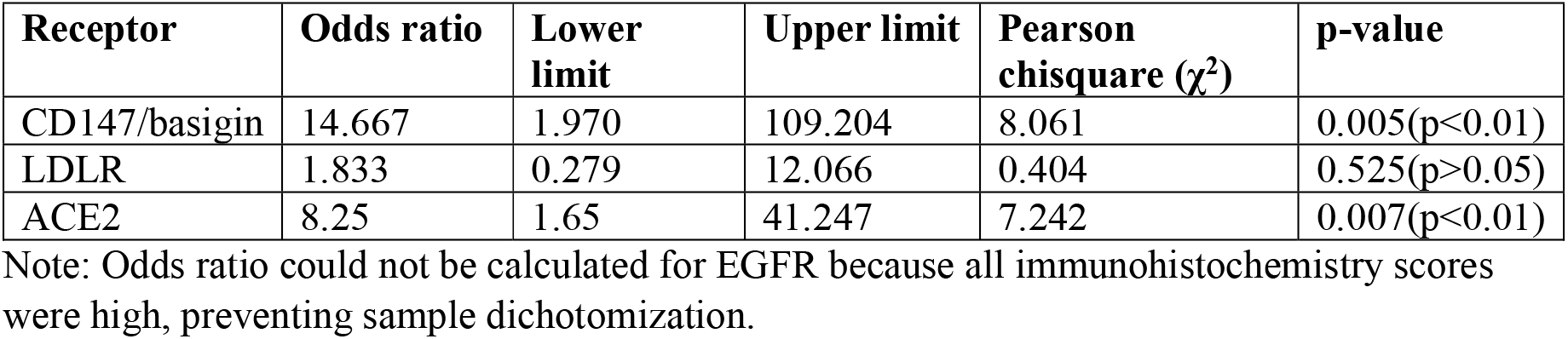
The association of receptor protein expression to COVID exposure risk obtained by immunohistochemistry.

| Receptor | Odds ratio | Lower limit | Upper limit | Pearson chisquare ($\chi^2$) | p-value |
| --- | --- | --- | --- | --- | --- |
| CD147/basigin | 14.667 | 1.970 | 109.204 | 8.061 | 0.005(p<0.01) |
| LDLR | 1.833 | 0.279 | 12.066 | 0.404 | 0.525(p>0.05) |
| ACE2 | 8.25 | 1.65 | 41.247 | 7.242 | 0.007(p<0.01) |

Note: Odds ratio could not be calculated for EGFR because all immunohistochemistry scores were high, preventing sample dichotomization.

### CD147/basigin, LDLR and EGFR bind to Spike protein, but not via the ACE2 receptor binding domain

To investigate whether the identified receptor proteins could be interacting with the SARS CoV-2 Spike protein, an initial *in silico* docking analysis was performed. Docking analysis suggested that CD147/Basigin, EGFR, and LDLR interact with the spike protein outside the receptor-binding domain (RBD). Interface residues were defined as residues with contact frequencies ≥ 0.60 during molecular dynamics simulations. In the spike-basigin complex (**Figure 4A**), the interfacial residues showed low backbone RMSD values and stabilized after 10 ns (**Figure 4B**), with a mean RMSD of 2.82 ± 0.10 Å, supporting a stable interface. Contact frequencies were calculated across all interaction classes, including hydrogen bonds, salt bridges, hydrophobic contacts, van der Waals interactions, π-cation interactions, π-stacking, and T-stacking. Residue-residue contacts with frequencies ≥ 0.85 are plotted in **Figure 4D** and mapped onto the initial docking pose in **Figure 4C**. Spike protein Tyr28 formed van der Waals contacts with basigin Asn98 and Ile99 with frequencies of 0.91 and 0.93, respectively, indicating that Tyr28 is a major contributor to complex stabilization.

**Figure 4:**
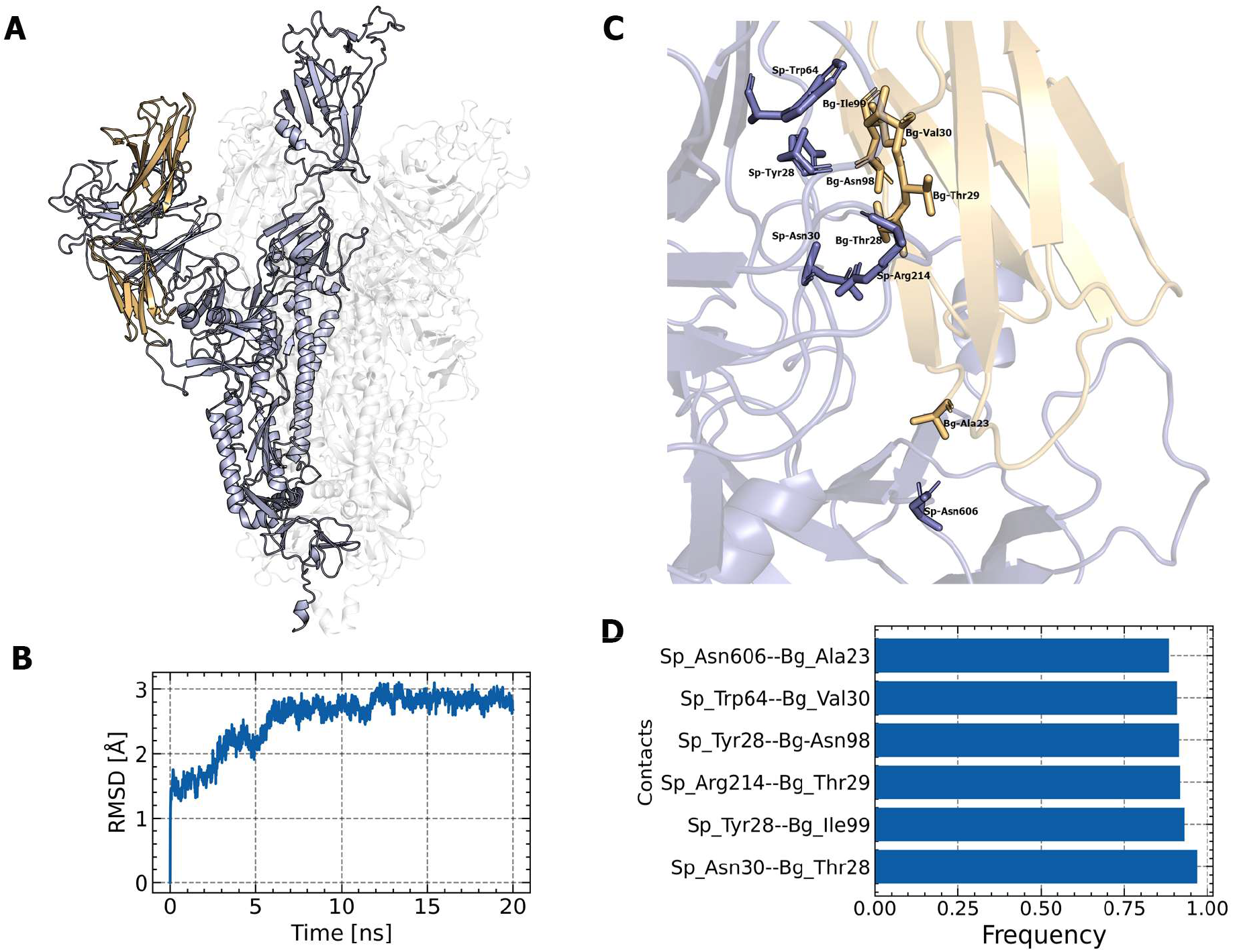
A) Initial docking conformation of the Spike protein chain B (purple) and Basigin (orange); B) RMSD of the backbone atoms of the interfacial residues; C) Interfacial residues with contact frequency ≥ 0.85 displayed on the initial docking structure; D) Contact frequency plot of the corresponding residues.

To capture structural diversity in EGFR (UniProt ID: P00533), a BLAST search against the Protein Data Bank (PDB) identified 11 EGFR structures with varying conformations and binding partners. These structures were clustered by backbone root-mean-square deviation (RMSD) using a 2 Å cutoff, yielding four distinct conformational states (**Figure S1**). Although 1IVO is structurally similar to 3NJP (backbone RMSD = 0.31 Å), it was retained to evaluate whether subtle conformational differences influence Spike recognition.

All selected EGFR conformations docked to a specific chain of the Spike glycoprotein (**Figure 5**); however, their binding poses differed substantially. During molecular dynamics simulations, each EGFR-Spike complex showed distinct interfacial dynamics. The Spike-3NJP and Spike-3QWQ complexes (**Figure S2 and S3**) exhibited low interfacial backbone RMSD values after 10 ns, with means of 2.00 ± 0.07 Å and 1.88 ± 0.10 Å, respectively (**Figures S2B and S3B**), suggesting stable interfaces. In contrast, the Spike-1IVO complex, despite its structural similarity to 3NJP, reached a higher plateau RMSD of 4.58 ± 0.10 Å between 5 and 10 ns (**Figure S6B**). The Spike-1NQL and Spike-5XWD complexes (**Figure S4 and S5**) also stabilized at higher RMSD values after 10 ns (3.50 ± 0.18 Å and 4.58 ± 0.10 Å, respectively), suggesting substantial rearrangement from the initial docked poses (**Figure S4A,B and S5A,B**).

**Figure 5:**
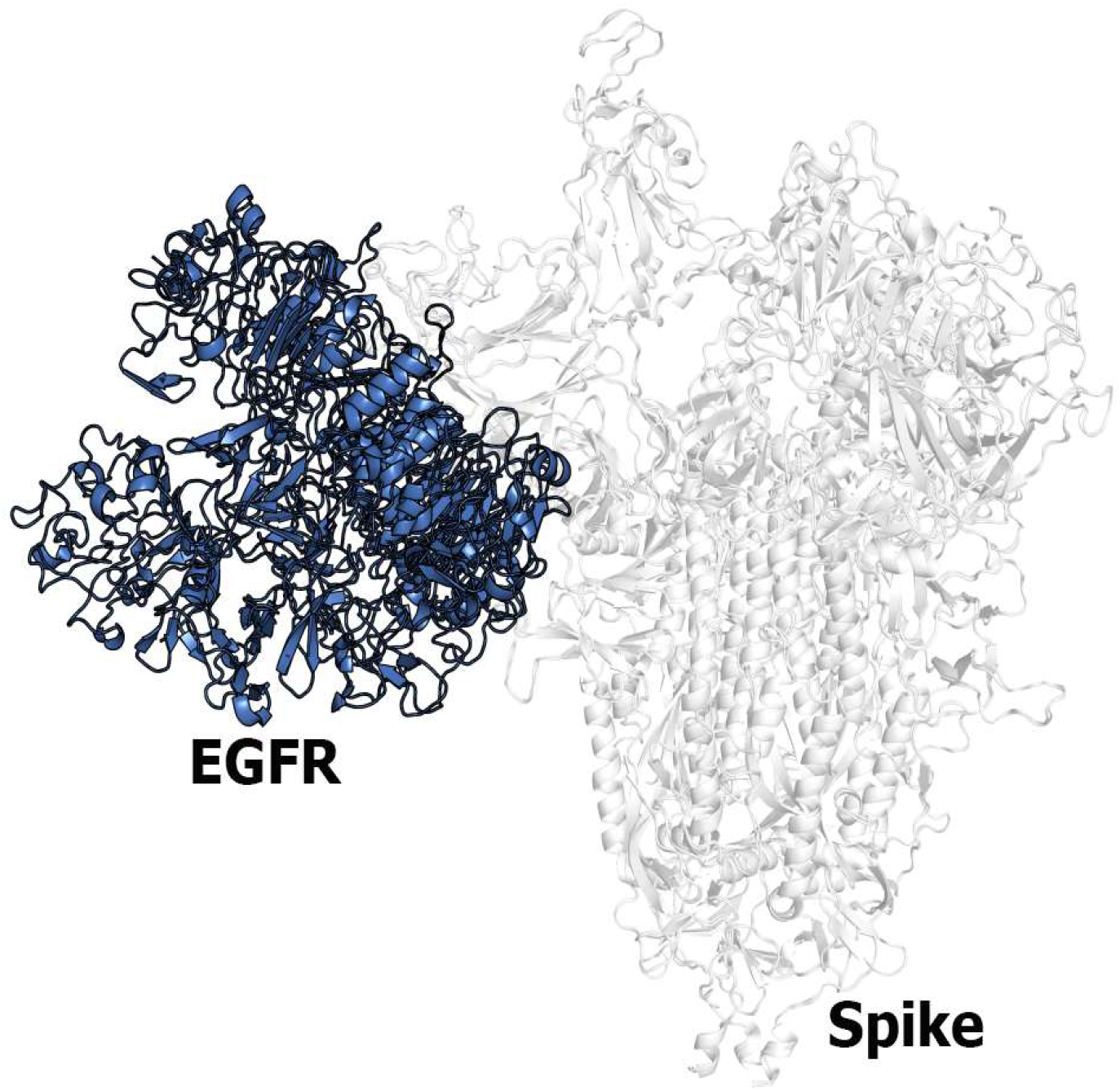
Top docking poses of different EGFR (Blue) conformations with spike glycoprotein (White).

Across the EGFR simulations, the contact frequency profiles (**Figure S2D-S5D**) and corresponding mapped interfaces (**Figure S2C-S5C**) identified Tyr28, Gly72, Arg214, Asp215, and His245 as recurrent Spike protein residues involved in EGFR binding. For EGFR, His159, Arg125, Pro349, Arg353, Asp355, Ser356, Phe357, and Gln384 were the principal residues contributing to Spike interactions across the sampled conformations.

The LDLR-Spike interface (**Figure 6**) remained stable throughout the simulation, with a mean interfacial backbone RMSD of 2.74 ± 0.13 Å after 10 ns. High-frequency contacts are mapped and quantified in **Figure 6C and 6D**.

**Figure 6:**
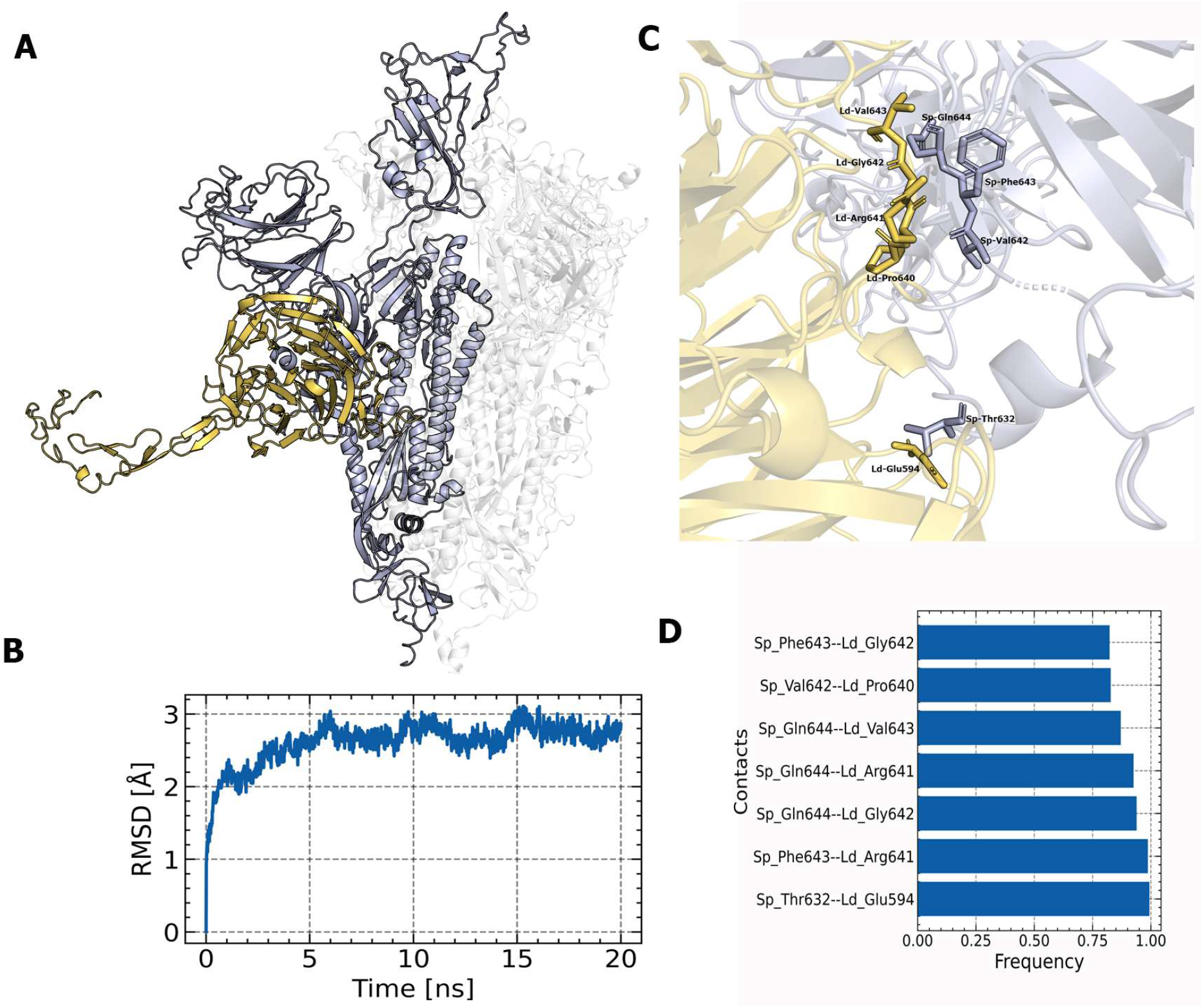
A) Initial docking conformation of the Spike protein chain B (purple) and LDLR (orange); B) RMSD of the backbone atoms of the interfacial residues; C) Interfacial residues with contact frequency ≥ 0.85 displayed on the initial docking structure (Sp: spike protein, Ld: LDLR); D) Contact frequency plot of the corresponding residues.

### Coimmunoprecipitation validation of Spike protein interaction with with basigin, LDLR and EGFR

To validate our *in-silico* analysis, immunoprecipitation reactions were set up. In all experiments, the protein of interest-basigin/CD147, EGFR and LDLR endogenously expressed in placental tissue lysates was immobilised on agarose beads via pre-coating using specific antibodies. These beads were subsequently incubated with 293T cell lysate overexpressing spike protein. Immunoblotting for spike protein revealed prominent bands in the lanes where placental lysate with antibody against the aforementioned proteins had been incubated with spike protein. (**Figure 7A, 7B, 7C**). However, very faint or no bands were seen in the lysate with bound protein incubated without antibody as control. Prominent bands were also observed in the tissue homogenate transfected with spike indicating that the expression of spike protein was specific to the transfected cells only.

**Figure 7:**
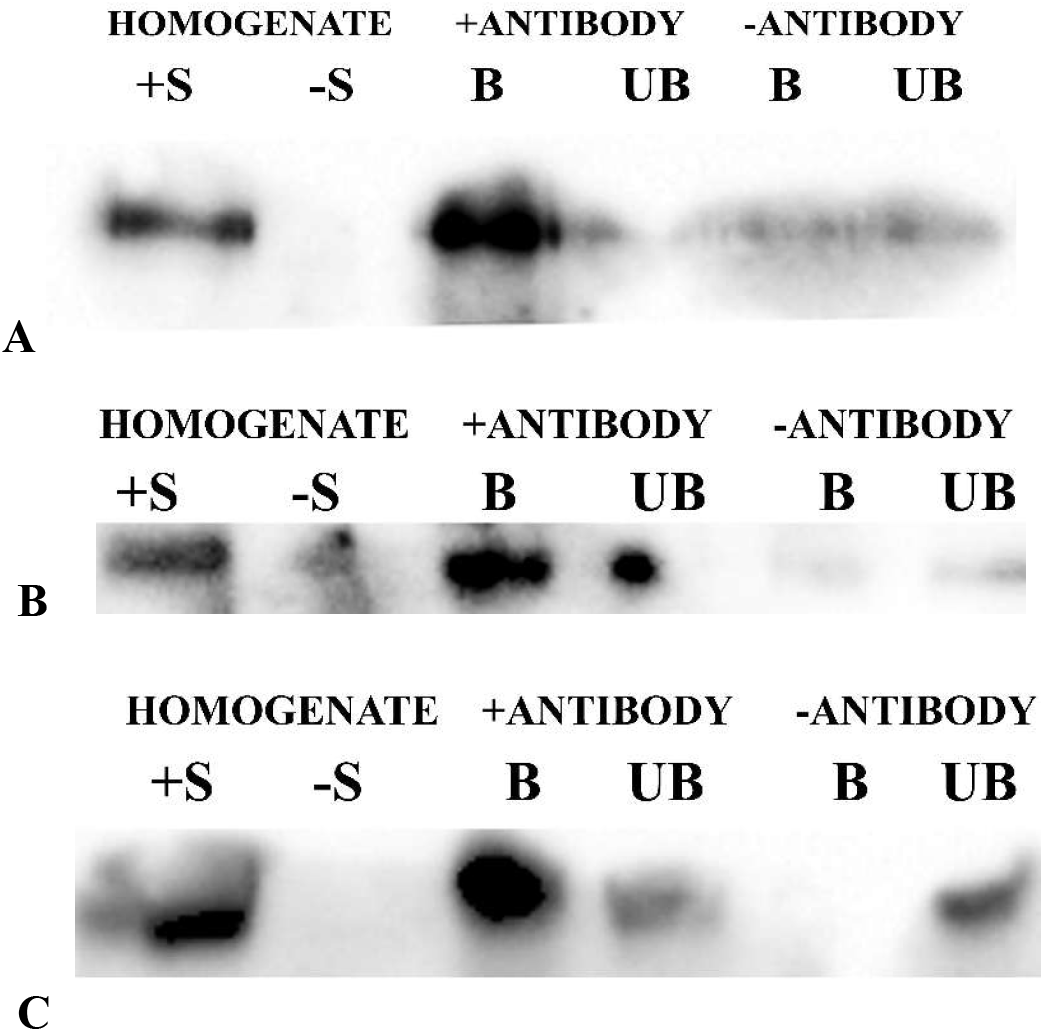
Co-immunoprecipitation of spike protein with CD147, EGFR and LDLR. In all cases the protein of interest (A) CD147, (B) EGFR and (c) LDLR was immobilised on agarose beads using specific antibody which was incubated with placental lysates expressing these proteins. Interaction with spike was assessed by incubation with spike expressing 293T cell lysates. Presence of spike was observed via immunoblotting for spike using specific antibody. +S indicates cell homogenate expressing spike, -S indicates control cell homogenate without spike, B indicates antibody bound fraction, UB the unbound fraction, +ANTIBODY: lysate with antibody for target receptor protein, -ANTIBODY: lysate without antibody for target receptor protein. Observed band size for spike is 110 kDa for A, B and C.

### *In-silico* saturation mutagenesis identifies receptor-specific Spike binding hotspots

To further evaluate the contribution of high-frequency interfacial residues, in-silico saturation mutagenesis was performed for the Spike-Basigin, Spike-LDLR, and Spike-EGFR complexes. The resulting heatmaps report the predicted change in binding free energy (ΔΔGbind) for each substitution, where positive values indicate mutations predicted to weaken binding, whereas negative values indicate substitutions predicted to strengthen binding relative to the corresponding wild-type residues (**Figure** 8).

**Figure 8:**
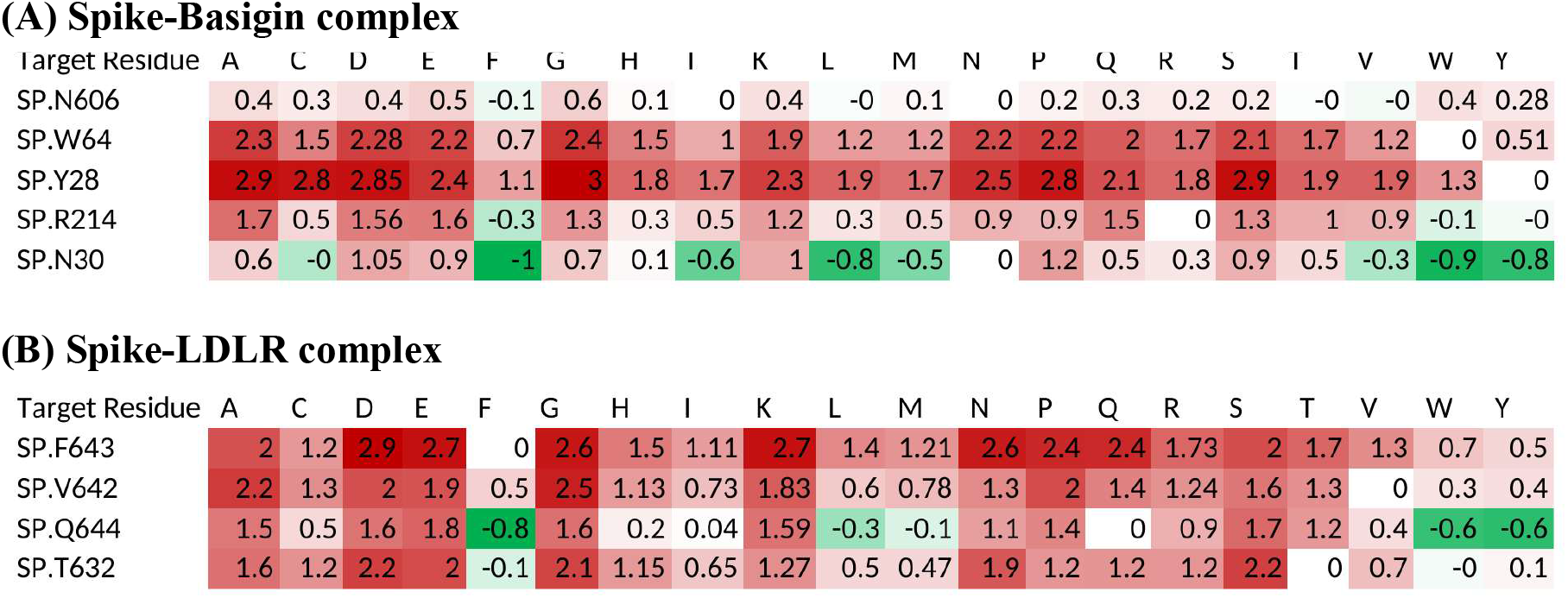
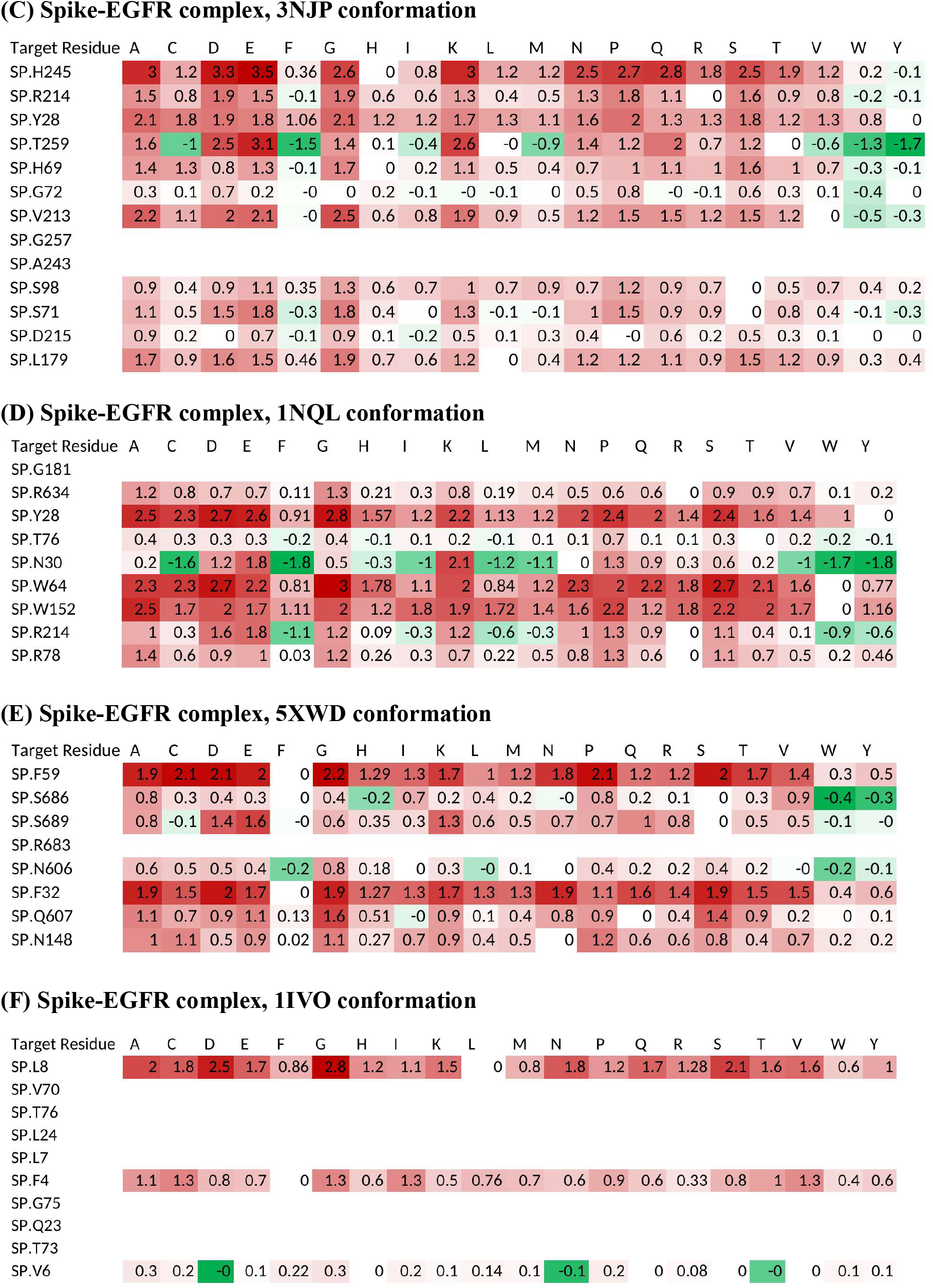
*In silico* saturation mutagenesis of Spike residues at the predicted receptor interfaces. Heatmaps show predicted mutation-induced changes in binding free energy (ΔΔGbind) for Spike residues in complex with (A) Basigin, (B) LDLR, and EGFR conformations (C) 3NJP, (D) 1NQL, (E) 5XWD, and (F) 1IVO. Red-shifted cells indicate substitutions predicted to decrease binding affinity, whereas green-shifted cells indicate substitutions predicted to increase binding affinity. The 3QWQ EGFR conformation did not return mutational-energy values and was therefore not included in this hotspot comparison.

For the Spike-Basigin complex, mutations at spike Trp64 and Tyr28 produced consistently positive ΔΔG bind values across most substitutions, indicating that these residues are major contributors to Basigin binding. In the Spike-LDLR complex, Phe643, Val642, and Thr632 showed similar destabilizing mutational profiles, supporting their role as LDLR-interaction hotspots.

For EGFR, the mutational analysis was performed across the available docked conformations. No mutational-energy values were obtained for the 3QWQ model; therefore, this conformation was excluded from the hotspot analysis. Across the remaining EGFR-Spike complexes, Leu8, Phe59, Phe32, Tyr28, Trp64, Trp152, His245, and Val213 emerged as key Spike residues, because mutations at these positions generally reduced the predicted binding affinity. These mutagenesis results are consistent with the contact-frequency analysis and further support a conformation-dependent Spike-EGFR recognition mechanism.

## Discussion

### Main findings

The study revealed a few noteworthy points with respect to potential co-factors which might be involved in vertical transmission of SARS-CoV-2 or at least placental susceptibility. While the spike protein primarily interacts with the ACE2 receptor in the RBD for entry, it interacts with EGFR, LDLR, and basigin/CD147, but not in the RBD or receptor binding domain. When their expression was assessed at mRNA and protein level, it was found that both basigin and LDLR had relatively higher expression in pregnant women with exposure risk to COVID than control group. In contrast, EGFR mRNA level was lower in the exposure risk group while, EGFR protein expression was only slightly high in the patients with exposure risk. Furthermore, *in vitro* validation confirmed the interaction of the spike protein with basigin/CD147, consistent with recent cryo-EM findings, and identified previously unreported interactions with EGFR and LDLR. We could also validate that Trp64, Tyr28 and Phe643, Val642, Thr632 are the key residues of spike which play crucial role in the interaction between spike-basigin and spike-LDLR complex respectively, through in-silico site-directed mutagenesis. Whereas, for spike-EGFR complex, Leu8, Phe59, Phe32, Tyr28, Trp64, Trp152, His245, and Val213 emerged as important spike protein residues. Therefore, the findings suggest the role of basigin, EGFR and LDLR as alternate interactors of spike in placenta besides the canonical receptor ACE2. It may be hypothesized that perhaps ACE2 serves as the primary receptor for the spike protein, specifically interacting with the RBD in the open conformation, while the aforementioned proteins may function as potential co-factors or spike-interacting partners.

### Interpretation

Our findings exhibit a trend of increased expression of CD147, LDLR and ACE2 both at transcription and translation level in placentas with high maternal COVID exposure risk. EGFR however deviate from this trend and show low mRNA expression and a slightly higher protein expression due to epidemiological exposure risk. Although, this change is not statistically significant, it indicates that maternal environmental risk of COVID may modulate placental receptor expression. This data is further supported by odds ratio analysis. The values for CD147 and ACE2 demonstrate statistically significant positive association between maternal exposure risk to SARS-CoV-2 and receptor expression at the protein level. Dichotomization could not be performed for EGFR protein expression which implies uniformly high expression irrespective of COVID exposure, reflecting its constitutive expression pattern rather than induced by viral exposure.

The computational study predicted that, CD147/basigin, EGFR and LDLR interact with spike protein outside the receptor binding domain/RBD suggesting their possible role as secondary receptors with respect to vertical transmission. The low RMSD values during simulation, contact frequency values together with binding hotspots through saturation mutagenesis suggests these interactions to be important. This reinforces the existing hypothesis that SARS-CoV-2 might interact with multiple receptors or co-factors besides its canonical receptor ACE2.

Importantly, these in-silico predictions were further confirmed by positive interactions of spike with basigin/CD147, EGFR and LDLR as spike co-precipitated with these proteins respectively. The absent or faint spike signal in the negative controls further rule out non-specific interactions. Therefore, these in-vitro experiments demonstrate that these three candidate proteins might act as alternate interacting partners of spike in the placenta.

### Strengths and limitations

The present study has its strengths as well as limitations. While current literature suggests the plausible role of CD147, EGFR and LDLR as cofactors for SARS-CoV-2 entry based on their expression in the placenta (14), there was no experimental evidence to confirm their interaction, especially for EGFR and LDLR. Early research was performed to computationally study the interaction between CD147 and spike. But their work was limited to the RBD and protonation state of the structures was not checked (63). Another interaction study of CD147 with spike protein revealed no direct interaction between them (64). Furthermore, while our work was in progress, a study on CD147 was published where the Cryo-EM analysis showed that CD147 binds to the SARS-CoV-2 spike RBD in “up” conformation. The loop connecting the two immunoglobulin-like domains of CD147 engages the RBD β6 strand and is stabilized by five hydrogen-bonding residue pairs: CD147 R54, E84, E92, Q100, and S112 with spike G413, K417, K424, Y489, and G447, respectively (65). This recently reported crystal structure identifies a CD147–Spike binding mode that is substantially different from our computationally predicted complex. Our ClusPro model predicts interactions primarily with other regions of Spike (e.g., residues Y28, N30, W64, R214 and N606). Such differences are expected in protein–protein docking, which predicts favorable binding poses but may not fully capture conformational changes associated with binding, glycosylation, membrane effects, or induced fit. Hence further *in vitro* studies are needed to confirm the exact site of interaction. However, our quest to find alternate interacting partners of spike or co-factors in placenta led to the finding that spike protein might interact with multiple receptors besides ACE2. Both computational and experimental observations confirm these interactions for the first time, especially for EGFR and LDLR, which is a novel revelation in the context of pleacental susceptibility to COVID. This strengthens the potential involvement of multiple receptors the virus might use for entry thus increasing cellular susceptibility. Also, with the revelation of alternate spike-interacting partners or co-factors, proper targeted therapeutic interventions could also be investigated.

However, from the current study we cannot confirm whether these interactions are direct or indirect. We can suggest that these proteins are a part of an interacting complex. Future experiments are required to establish the nature of these interactions. Whether these interactors have the potential as alternate receptors, could be validated by studying SARS-CoV-2 pseudovirus entry in a placental trophoblast cell line with knock-down of these spike-interacting proteins individually. Also, among the 12 interactors found through interactome analysis this study is limited to three receptors only. The study can be continued to explore if the other receptors played any noteworthy role in vertical transmission of SARS-CoV-2.

### Conclusions

In conclusion we can state that our findings provide a novel insight into non-canonical interacting partners in the vertical transmission of SARS-CoV-2. Through expression profiling, computational modelling and experimental validation, this study identifies potential multiple co-factors of spike along-with ACE2 in placenta which increases the susceptibility to virus attachment. Finally, our work provides a foundation for future research into molecular mechanism of placental SARS-CoV-2 infection and identifies potential secondary co-factors or attachment factors as therapeutic targets to reduce the chances of mother-to-foetus virus transmission.

## Author contributions

Sulagna Mandal: Methodology, Validation, Formal analysis, Investigation, data curation, Writing-original, review and editing,

Satyabrata Majumder: Methodology, Validation, Formal analysis, Investigation, Writing-original, review and Editing

Suman Kumar Nandy: Formal analysis, Investigation

Ruchira Mukherjee: Formal analysis

Kalyan Giri: Writing - Review & Editing, Supervision

Aparna Mukhopadhyay: Conceptualisation, Writing - Review & Editing, Supervision, Project administration

## Ethical Approval

This work involves human samples. Ethical clearance was obtained from Calcutta Medical College Hospital, Kolkata and Calcutta National Medical College Hospital, Kolkata vide ethical certificate: MC/KOL/IEC/NON_SPON/1290/03/2022 dated 16/03/2022 and EC-CNMC/2024/403 dated 04/05/2024. Signed informed consents were obtained from all patients participating in the study. No animals were used.

## Data Availability

Much of the supporting data is uploaded as supplementary information. The docked structures are uploaded in in zenodo via DOI: https://doi.org/10.5281/zenodo.23157295, https://doi.org/10.5281/zenodo.23157187, https://doi.org/10.5281/zenodo.23156901 and can be made available upon request.

## Acknowledgements

Acknowledgement is due to Prof. Amlan Ghosh for his help in histopathological studies. An acknowledgement is extended towards Ms. Payal Bairagi for her role as second observer in the immunohistochemistry scoring.

## Declaration on use of AI

AI was used for paraphrasing and English corrections. For such purpose Gemini, Claude, ChatGPT was used.

